# Composite receptive fields in the mouse auditory cortex

**DOI:** 10.1101/2021.10.13.464267

**Authors:** Sihao Lu, Mark Steadman, Grace W. Y. Ang, Andrei S. Kozlov

## Abstract

A central question in sensory neuroscience is how neurons represent complex natural stimuli. This process involves multiple steps of feature extraction to obtain a condensed, categorical representation useful for classification and behavior. It has previously been shown that central auditory neurons in the starling have composite receptive fields composed of multiple features when probed with conspecific songs. Whether this property is an idiosyncratic characteristic of songbirds, a group of highly specialized vocal learners, or a generic characteristic of central auditory systems in different animals is, however, unknown. To address this question, we have recorded responses from auditory cortical neurons in mice, and characterized their receptive fields using mouse ultrasonic vocalizations (USVs) as a natural and ethologically relevant stimulus and pitch-shifted starling songs as a natural but ethologically irrelevant control stimulus. We have found that auditory cortical neurons in the mouse display composite receptive fields with multiple excitatory and inhibitory subunits. Moreover, this was the case with either the conspecific or the heterospecific vocalizations. We then trained the sparse filtering algorithm on both classes of natural stimuli to obtain statistically optimal features, and compared the natural and artificial features using UMAP, a dimensionality-reduction algorithm previously used to analyze mouse USVs and birdsongs. We have found that the receptive-field features obtained with the mouse USVs and those obtained with the pitch-shifted starling songs clustered together, as did the sparse-filtering features. However, the natural and artificial receptive-field features clustered mostly separately. These results indicate that composite receptive fields are likely a generic property of central auditory systems in different classes of animals. They further suggest that the quadratic receptive-field features of the mouse auditory cortical neurons are natural-stimulus invariant.

## 1 Introduction

Since it was first described in single optic nerve fibres of the frog [Hartline, 1938], the concept of the receptive field has proved an important tool to characterize neuronal functionality throughout the central nervous system. It refers to a region in sensory space that can modulate a neuron’s activity. In the visual system, for example, this could correspond to a specific range of coordinates within the visual field within which brightness can modulate neuronal firing rate. In the auditory system, it might correspond to a particular range of frequencies and intensities.

One pervasive issue in characterizing neuronal receptive fields is the typically high dimensionality of the stimulus. To fully characterize the influence of each dimension on neuronal output, one would need to use impractically large stimulus sets and record responses over an unrealistically extended period. As a result, synthetic stimuli whereby only a small number of parameters of interest are manipulated have been typically used. A common example of this approach in the auditory system is the use of pure tones, played at a range of frequencies and intensities. Measuring the neuronal firing rate in response to each of these tones yields the pure-tone receptive field, frequency response area or frequency tuning curve. Early experiments using pure tones yielded important insights into the frequency-to-place mapping that occurs in the cochlea and is maintained throughout the auditory system.

Subsequent experiments using more complex sounds have indicated highly non-linear response properties, meaning that it is difficult to predict a neuron’s response to more complex stimuli by probing its response to individual pure tones [Machens et al., 2004]. Therefore, it seems that the estimated receptive field is to a large degree determined by the stimulus chosen to measure it. Given that one of the primary goals of sensory neuroscience is to understand the relationship between stimulus and behavior, it is important to understand how neurons respond to behaviorally relevant, natural stimuli. In vocalizing animals (including humans), a strong candidate for such stimuli would be conspecific vocalizations.

Historically, there have been a number of barriers to successfully estimating receptive fields using complex, natural sounds. Firstly, traditional methods to characterize neuronal input-output relationships have been based on a number of assumptions that do not hold true in the case of natural stimuli. The spike-triggered average (STA, or reverse-correlation method [de Boer and Kuyper, 1968]), for example, requires that the stimuli are spherically symmetric (i.e., uniformly randomly sampled). In the auditory domain, such a stimulus is known as white noise, to which the auditory cortex appears to adapt leading to relatively weak responses over time [King et al., 2018]. One approach to overcome the need to use white noise is to effectively “whiten” the stimulus before carrying out reverse-correlation analysis by compensating for intrinsic stimulus correlations [Theunissen et al., 2000]. However, neurons in the neocortex can receive inputs from thousands of excitatory and inhibitory synapses of diverse origin simultaneously, so it seems likely that a receptive field characterized by a single spectro-temporal “feature” would often be insufficient to fully describe the neuronal input-output relationship. The spike-triggered covariance (STC) is one method that yields a receptive field comprising multiple “features” (each corresponding to a unique combination of stimulus dimensions that appears to modulate neuronal activity), but like reverse-correlation, this method is also beset by a number of assumptions that do not hold true in the case of natural stimuli.

More recently, information-theoretic approaches have been proposed to overcome the challenges described above and enable composite receptive field estimation from responses to natural sounds. The Maximally Informative Dimensions (MID) method [Sharpee et al., 2004] facilitates receptive-field estimation by searching for a feature that maximizes the mutual information between the stimulus and the spikes. Whilst this approach can be extended to estimate receptive fields with multiple features (like the STC [Rowekamp and Sharpee, 2011]), it is computationally intensive and limited by the impractical amount of physiological data required to estimate receptive fields comprising more than a small number of features.

An alternative approach is the Maximum Noise Entropy method [Fitzgerald et al., 2011]. This method optimizes a model that maps the stimulus onto a logistic function, the output of which represents the probability of a spike. The parameters of this model are constrained by stimulus-response correlations up to the second order (i.e., corresponding to overall firing rate, spike-triggered average, and spike-triggered covariance), but it is otherwise as random and hence unbiased as possible. This method has been applied to neurons in the auditory forebrain of songbirds [Kozlov and Gentner, 2016], which demonstrated that these neurons had composite receptive fields composed of multiple excitatory and inhibitory features.

The songbird auditory forebrain (specifically the caudo-medial nidopallium, NCM) appears to play a particularly important role in processing birdsong and thus may have functionally diverged from the mammalian auditory cortex. More recently, Atencio and Sharpee [2017] applied the MNE model to characterize the responses of neurons in the cat auditory cortex, in response to dynamic moving ripples and ripple noise. That study also found that individual auditory neurons have multidimensional receptive fields, albeit in response to artificial sounds. Another study, in the ferret auditory cortex, discovered multiple excitatory and inhibitory features in single neurons driven by an ensemble of natural sounds comprising environmental sounds, ferret vocalizations, and human speech [Harper et al., 2016]. An emerging theme from these studies is that central auditory neurons have composite receptive fields, and that their properties may differ depending on whether they are probed with artificial or natural sounds.

In this study, we wanted to answer the following questions: When auditory cortical neurons in the mouse, a powerful model organism in neuroscience, respond to ethologically relevant mouse vocalizations (USVs), what is the dimensionality of the feature space? Does it differ depending on whether neurons are stimulated with ethologically relevant sounds (mouse USVs) or with natural, but ethologically irrelevant complex sounds (pitch-shifted birdsongs)? Finally, how do these features compare to those learned by an artificial neural network optimized for sparse representation?

## 2 Materials and Methods

### 2.1 Surgical preparation

All procedures were carried out under the terms and conditions of licences issued by the UK Home Office under the Animals (Scientific Procedures) Act 1986. Extracellular recordings were made in the auditory cortex of adult female C57BL/6 mice (N=25, aged 5 to 27 weeks). Animals were anaesthetized using a mixture of fentanyl, midazolam and medetomidine (0.05, 5 and 0.5 mg/kg respectively). A midline incision was made over the dorsal surface of the cranium and the left temporalis muscle was partially resected. The location on the skull over the left auditory cortex was identified using a rostro-caudal coordinate of 70% bregma-lambda and a dorso-ventral coordinate of bregma -2.2 mm (the lateral coordinate being determined by the surface of the skull). A steel headplate comprising a bent piece of flat bar (approximately 5 mm x 30 mm) was attached to the dorsal surface of the skull using a combination of tissue adhesive (Histoacryl) and dental cement. This was subsequently used in combination with a magnetic stand to secure the animal in place for the remainder of the surgery and recordings.

A small (∅=2mm) craniotomy was made over the auditory cortex using a dental drill and burr. For recordings made using multi-shank tetrode arrays, an incision was made in the dura using a small needle, through which the probes were later inserted. For single shank polytrode recordings, the dura was left intact prior to insertion. The surface of the brain was protected from desiccation by regular application of warmed saline solution or agar. A second craniotomy was made approximately over the contralateral motor cortex, such that a machine screw (M1 × 2 mm) could be secured into the bone to provide a reference signal. The animal was then transferred to an acoustic chamber, where the head was again secured by means of the headpost and a magnetic stand and body temperature was maintained throughout the duration of the experiment using activated Deltaphase isothermal pads.

### 2.2 Electrophysiological recording

Extracellular signals were recorded using silicon multi-electrode probes (Neuronexus). Two styles of probe were used to obtain the data presented here: a 4-shank, 4 by 2 tetrode array and a single-shank, 32-channel polytrode. The tetrode array facilitated simultaneous recordings at sites spanning the tonotopic map, whereas the polytrode provided greater robustness to electrode drift. For both tetrodes and polytrodes, the conductive site areas were 177*µm*^2^. Signals were amplified and digitized at a sample rate of 40 kHz using a Plexon Omniplex Neural Recording Data Acquisition System (Plexon Inc.). A subset of the recordings were acquired through a Neuronexus Smartbox Pro (Neuronexus) at 30 kHz.

### 2.3 Stimuli

Mouse ultrasonic vocalizations were obtained from mouseTube [Torquet et al., 2016], an annotated database of ultrasonic mouse calls. The database was searched for vocalizations produced by male C57BL/6 mice. These were downloaded and individual song bouts were extracted manually using Audacity. The extracted bouts were ramped on and off using a 10-ms cosine squared ramp and concatenated into a single audio file with a sample rate of 250 kHz. Two versions of the USV stimulus were produced. The first had a total duration of 3 minutes and 30 seconds. The second version was edited such that periods of silence or movement noise between song bouts were shortened. This stimulus had a total duration of 2 minutes.

In addition, a stimulus based on starling vocalizations was also produced. Starling song recordings were taken from the database produced by Kozlov and Gentner [2016]. Briefly, spontaneously produced songs of adult male European starlings (*Sturnus vulgaris*) were recorded in a sound attenuation chamber at 44.1 kHz. These were then upsampled to 250 kHz and pitch-shifted up by a factor of 10 to better match the power spectra of the mouse vocalizations. Similar to the mouse vocalizations, two versions of the starling stimuli were used: the first had a total duration of 3 minutes 30 seconds and the second was more condensed, with periods of silence removed and had a total duration of 2 minutes. All audio files were adjusted such that the RMS intensities were matched. An example of both these stimuli can be seen in the top two panels of Figure 1.

**Figure 1:**
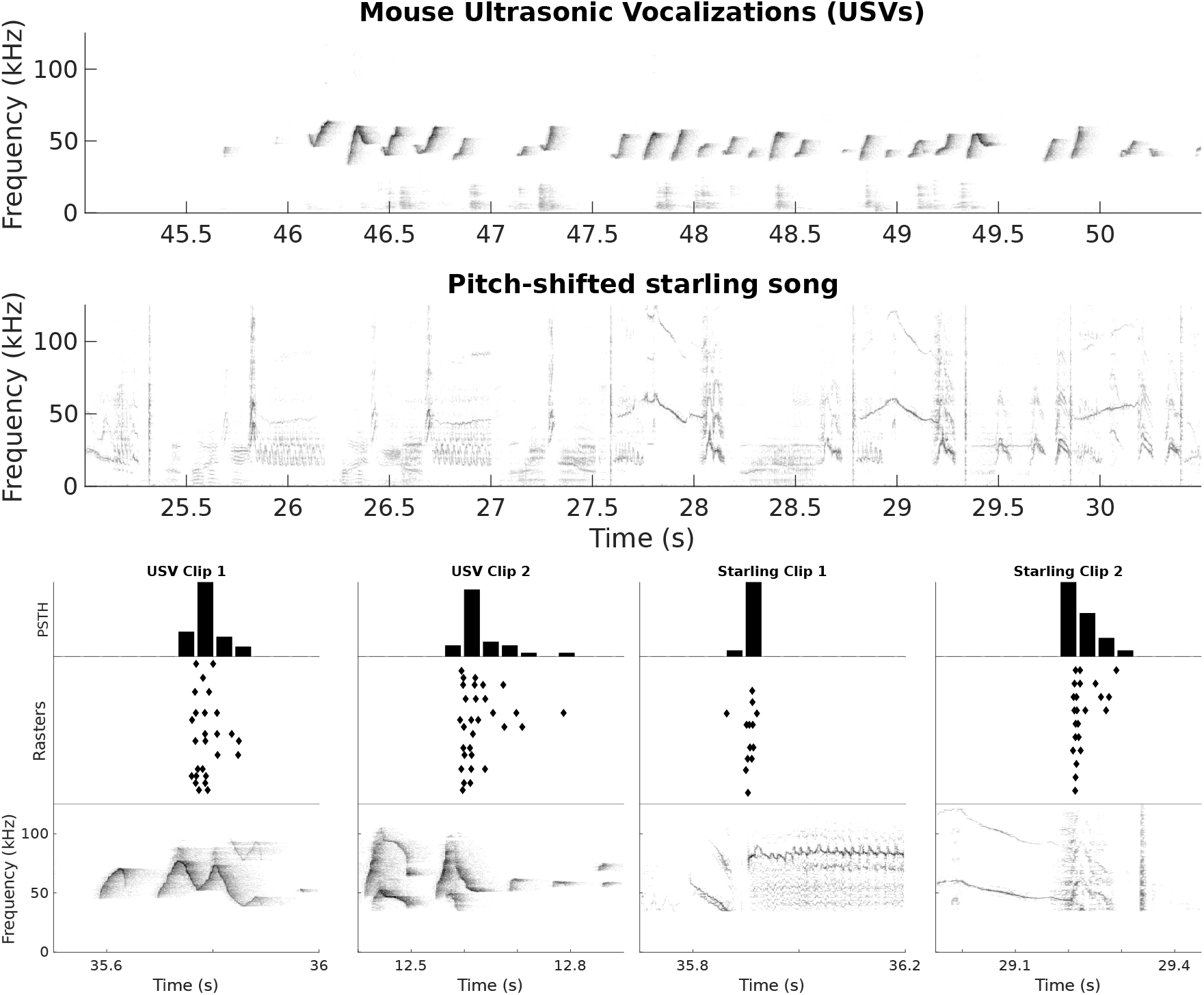
Examples of stimuli and responses. The top panel shows a spectrogram of a 5.5-second fragment of a mouse ultrasonic vocalization stimulus. The second panel shows a spectrogram of a 5.5-second fragment of a starling song, pitch-shifted up by a factor of 10 to better match the overall spectrum of the mouse USVs. The bottom panel shows short, 0.5-second fragments of both the USV and starling stimuli alongside the corresponding spiking response. These plots comprise the spectrogram of the stimulus, a raster plot of spiking responses whereby each dot represents a single spike, the x-coordinate corresponding to the time and the y-coordinate corresponding to the stimulus repetition number. Above the raster is a post-stimulus time histogram (PSTH).

Stimuli were presented for a minimum of 10 repetitions, up to a maximum of 20 repetitions, using an Avisoft UltraSoundGate Player 116 (Avisoft Bioacoustics) connected to a PC. In all cases, the sound level was adjusted such that the peak intensity did not exceed 85 dB SPL. Playback was monitored using an UltraSoundGate 166H coupled with an Avisoft-Bioacoustics CM16/CMPA condenser microphone connected to the same PC. Signal intensity was monitored in dB SPL by calibrating the system using a calibrated 40 kHz reference signal generator (Avisoft Bioacoustics). In order to synchronize neural responses and the presented stimuli, a digital output from the Player 116 was connected to a digital input in the data acquisition system, which triggered an event at the onset of stimulus playback.

### 2.4 Spike sorting

Spikes were sorted offline either using a semi-automated principle components clustering approach implemented in Plexon Offline Sorter (Version 4.4.0) or Kilosort3 [Steinmetz et al., 2021]. For the Plexon Offline Sorter, spike events were detected by setting a threshold close to the noise floor on each electrode. Detected events were then sorted and classified as either single- or multi-unit. The criteria for classifying a unit as single were that there was a locally consistent spike shape where the distribution of peak amplitudes was distinct from the spike detection threshold. Clustering was performed using the built-in k-means algorithm, the results of which were inspected and corrected visually using 3-D visualizations of a number of parameters including the first three principle components, maxima, minima and overall energy of the detected spike events. Since extracellular responses at a single recording location were recorded over time period of up to 2 hours, some recordings exhibited electrode drift, whereby spike shapes changed gradually over time. Therefore, all sorting was carried out using a 3-D clusters vs. time visualization. Spikes extracted using the automatic spike sorting algorithm Kilosort3 were manually curated with Phy [Rossant and Harris, 2013]. Units were considered well-isolated single units if they showed clear refractory periods in their autocorrelogram and had fewer than 1% spikes within a 1-ms interspike interval.

### 2.5 Receptive field characterisation

The receptive fields of single units were characterized using the Maximum Noise Entropy (MNE) method [Fitzgerald et al., 2011]. Briefly, this method comprises fitting a model that describes the probability of a spike as a function of a stimulus, **s**, as *P* (*spike*|**s**) = (1 + *exp*(*a* + **s***h* + **s**^*T*^ *J* **s**)^*−*1^, where *a, h* & *J* are parameters optimized via gradient descent. These parameters are determined such that the predicted firing rate, spike-triggered average and spike-triggered covariance match those in the observed data. The choice of how to represent the stimulus can have a profound effect on the success of the model. For auditory data, the most straightforward representation would be the sample values of the acoustic waveform. However, this would yield a stimulus representation with intractably large dimensionality. It is therefore common to represent an auditory stimulus in terms of a spectrogram. The frequency limits of the spectrogram were determined through a combination of bioacoustical analysis of the stimulus set and published information on the behavioral audiogram of C57BL/6 mice.

Bioacoustical analysis of the USV stimuli was carried out using Avisoft SAS Lab Pro (Version 5.2.09). Spectrograms of the the entire ultrasonic vocalizations corpus were generated using a 512-point Hann window and 50% overlap. Individual USV syllables were detected and analyzed using a semi-automatic thresholding process, whereby a threshold of -50 dB to -45 dB relative to the maximum intensity (with a hold time of 10 ms) was used. This method would occasionally erroneously detect broadband, movement-related sounds, or else combine elements occurring in rapid succession, so detected elements were manually inspected and corrected where necessary. Whistle contours were then extracted by measuring the frequency of the highest intensity at intervals of 5 ms throughout each element. This analysis yielded a database of 1616 individual USV syllables with a mean duration of 70.6 ms (*σ*=55 ms) and overall average frequency of 55.4 kHz. In order to determine a suitable frequency range that covers the majority of the vocalization spectra, the minimum and maximum frequency (fmin and fmax, respectively) of each whistle contour were extracted. A lower frequency bound was set to 2 standard deviations below the mean fmin (27.3 kHz). An upper bound was set to 3 standard deviations above the mean fmax (100.6 kHz). Similarly, a maximum lag was set to 2 standard deviations above the mean duration (*µ* = 70.6 ms, *σ* = 55 ms), and rounded to the nearest bin size (180 ms). Since the behavioral audiogram of the mouse extends to lower frequencies, a small number of broader, low frequency bins were also included. Spectrograms were generated from the audio files using the Matlab function spectrogram. This was calculated using a 1024-point Hann window and a 50% overlap, yielding a representation with frequency bins separated by 244.1 Hz. A set of edge frequencies were defined, within which frequency bins were averaged. These were logarithmically spaced such that the resulting representation had 6 low frequency bins spanning from 18.1 kHz to 27.3 kHz and 26 high frequency bins spanning 27.3 kHz to 100.6 kHz. A temporal bin size of 15 ms was used. Columns of the spectrogram corresponding to times falling within each 15 ms bin were also averaged together. The intensity of the resulting spectrogram was then converted to a dB scale.

Since for each temporal bin the spectro-temporal receptive field (STRF) would comprise the spectrogram values at the same time and also during the preceding 180 ms, the spectrogram was concatenated with temporally lagged version of itself, so each row now comprised the dB intensity values in each of the 32 spectral bins, followed by the intensity values on the spectral bins at the previous timesteps, up to the maximum lag. This resulted in an *MxN* stimulus matrix, where *M* is the total number of time steps and *N* is equal to the number of spectral bins multiplied by the number of timesteps in the STRF, in this case equal to 384 (32 spectral bin multiplied by 12 timesteps). For the response, a post-stimulus time histogram (PSTH) was calculated using the same temporal bins and smoothed using a 225-ms Hanning window.

The MNE model for each unit was trained with 80% of the stimulus-response data. The remaining 20% of the data were used to test the model. The training data were randomly extracted from the full recording without replacement to minimize the effects of any non-stationarity of the data. After training the MNE model for each unit, the model was used to predict a response to the test stimulus. The correlation coefficient between the model predictions and the recorded neural activity was calculated to assess the performance of the model.

Neural activity of a single unit is not perfectly correlated between repeated presentations of the stimulus; this affects how the performance of a model based on its correlation coefficient with recorded data is interpreted. To account for this inherent variability in the neural activity, we estimate the expected correlation coefficient between the responses to repeated presentations of the same stimulus in the test set [Hsu et al., 2004, Touryan et al., 2005]. The expected correlation coefficient 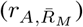 is defined as the correlation between the true firing rate, *A*, and the firing rate 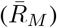 as measured by averaging over *M* repeats. As detailed in Hsu et al. [2004], we can calculate 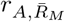 using the following equation:

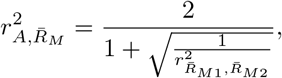

where 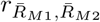 is the correlation between 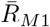 and 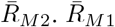 is the firing rate estimated by averaging over *M/*2 repetitions and 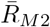 is the firing rate estimated by averaging over the other *M/*2 repetitions of the neural response. The correlation coefficients between the model prediction and the test set were then scaled by the expected correlation of the test set.

### 2.6 Feature estimation

In order to construct a compact representation of both the starling songs and mouse USVs, we used sparse filtering [Ngiam et al., 2011] to learn a set of basis features subject to the following constraints: populations sparsity, lifetime sparsity and high dispersal. Population sparsity specifies that only a small subset of features should be active at a given time, whilst lifetime sparsity specifies that each individual feature should be minimally active. High dispersal means that all the features have a similar amount of activity. One can compare this property with divisive normalization in neural circuits. These three constraints are typically observed in the neural activity in the cortex. How these constraints are applied is specified in [Ngiam et al., 2011]. Briefly, sparse filtering aims to find features based on their activity defined as *F* = *W * X* where *F* is the activity matrix which gives the activity of the features in *W* for each of the examples given in *X*. Each row *w*_*i*_ of *W* is one feature and each column *x*_*j*_ of *X* is one example. Thus each element *f*_*ij*_ in *F* defines the activity of feature *w*_*i*_ for example *x*_*j*_. The algorithm minimizes a cost function which takes the activity matrix as input by iteratively changing the features in *W* until an optimal set of features is found. Sparse filtering requires a single hyperparameter, the number of features to compute. For our analyses, we set the number of features to 128.

### 2.7 Latent space projection

To compare (1) the receptive-field features obtained from the neural activity with (2) the features estimated from the stimulus using the sparse filtering method as well as with (3) syllables segmented from the raw stimulus, we project the features into a latent space using the UMAP algorithm [McInnes et al., 2018]. The UMAP algorithm finds low-dimensional representations and has been shown to distinguish successfully between vocalizations of various animal species [Sainburg et al., 2020].

The syllables used in the projection were manually segmented from the stimulus, which resulted in 716 USV syllables and 1010 starling syllables. From this set of syllables we excluded any USV syllables shorter than 90 ms and any starling syllables shorter than 150 ms. This resulted in a final set of 552 USV syllables and 388 starling syllables. Spectrograms of the individual syllables were created using the same parameters as described in 2.5. Both the MNE features and the features estimated with sparse filtering were zero-padded to match the size of the spectrogram of the longest individual syllable. In addition to the segmented syllables and features obtained from MNE and sparse filtering, we added randomly generated noise in the form of spectrograms with the same number of frequency bins as the other inputs and a random time bin, up to the longest segmented syllable. The distribution of values in the noise spectrograms was chosen to be uniform.

Prior to approximating the latent space, the modulation power spectra of all the inputs were calculated by taking the two-dimensional Fourier transform of the spectrograms.

## 3 Results

### 3.1 Mouse auditory cortical neurons have composite receptive fields

Neuronal spiking activity in response to acoustic stimulation with mouse ultrasonic vocalizations was recorded from a total of 74 single units in the left auditory cortices of 25 anaesthetized mice. We also recorded responses to pitch-shifted starling song in a subset of these units (n = 54). Figure 1 shows examples of the recorded spikes during stimulation of both mouse USVs and pitch-shifted starling songs.

The receptive fields produced by the MNE method model yield a single linear feature (analogous to the spike-triggered average) and a number of quadratic features (analogous to the spike-triggered covariance). Figure 2 shows examples of the three most significant quadratic excitatory and inhibitory features of a single unit alongside their normalized eigenvalues indicating their relative contribution to the composite receptive field. A previously reported statistical method [Kozlov and Gentner, 2016] was employed to determine a number of significant excitatory and inhibitory features for each unit (Figure 2B). The numbers of excitatory and inhibitory features for all the units are summarized in Figure 3A. On average, the composite receptive fields comprised more significant inhibitory features (*µ*=8.17, *σ*=2.29) than excitatory ones (*µ*=6.85, *σ*=3.71). A two-sample Kolmogorov-Smirnov test confirmed that this difference was statistically significant (*P <* 0.001).

**Figure 2:**
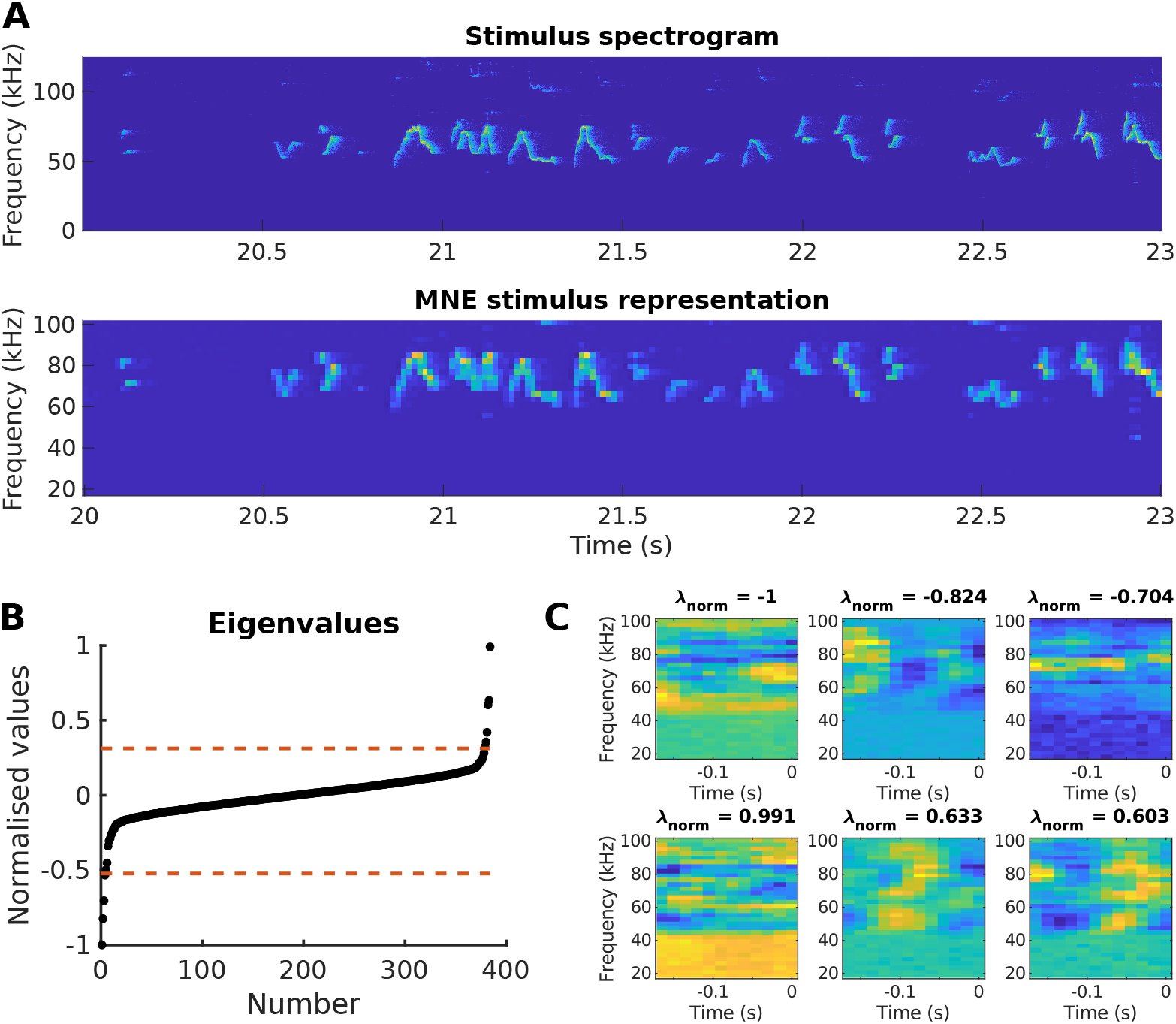
**(A)** The top panel shows a spectrotemporal representation of a section of the stimulus. The second panel shows the same segment after reducing the temporal resolution and logarithmically spacing the frequency bins; this representation was used to train the maximum noise entropy model. **(B)** The eigenvalues of all of the composite features along with the significance thresholds (dotted line). The features associated with the eigenvalues beyond the significance bounds are considered to be significant. **(C)** The three most significant excitatory (top row) and inhibitory (bottom row) features with their associated eigenvalues (*λ*_*norm*_). The eigenvalue determines the feature’s contribution to the composite receptive field.

**Figure 3:**
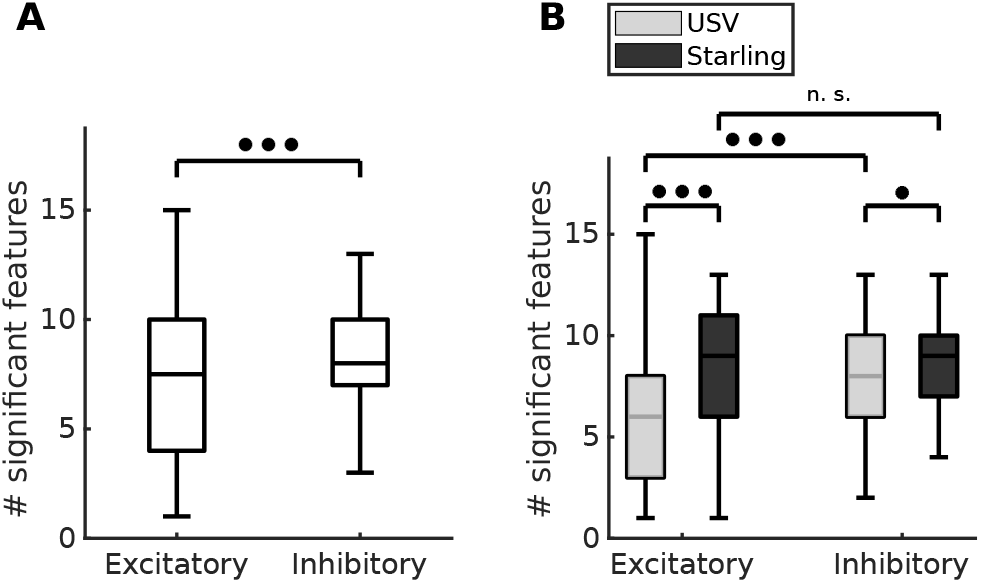
**(A)** Number of significant excitatory (*µ* = 6.85, *σ* = 3.71) and inhibitory features (*µ* = 8.17, *σ* = 2.29) for each of the receptive fields estimated. \*\*\**P <* 0.001 (two-sample Kolmogorov-Smirnov test). **(B)** Comparison between the number of significant features when the MNE model is trained with responses to USVs (*n* = 74) versus responses to pitch-shifted starling songs (*n* = 54). \*\*\**P <* 0.001 (two-sample Kolmogorov-Smirnov test), \**P <* 0.05 (two-sample Kolmogorov-Smirnov test). Every neuron stimulated with the starling songs was also stimulated with USVs.

### 3.2 Receptive fields estimated from con- and heterospecific vocalizations

In addition to mouse ultrasonic vocalizations, responses to pitch-shifted starling song of matched duration were recorded in 54 of the single units to understand better the importance of ethological relevance of the stimulus to estimating receptive fields. The receptive fields of these units were estimated using the two stimuli separately.

A comparison between the number of significant features estimated from conspecific and heterospecific vocalizations is shown in Figure 3B. We show that for the conspecific stimulus, the difference between the number of significant excitatory features (*µ* = 5.86, *σ* = 3.59) and the number of significant inhibitory features (*µ* = 7.70, *σ* = 2.31) is significant (*P <* 0.001, two-sample Kolmogorov-Smirnov test). On the other hand, there is no significant difference between the number of excitatory features (*µ* = 8.20, *σ* = 3.47) and the number of inhibitory features (*µ* = 8.81, *σ* = 2.11) estimated from the heterospecific stimulus (*P* = 0.28, two-sample Kolmogorov-Smirnov test). In addition, we show that the number of significant excitatory features differs between models trained using the conspecific stimulus and those trained using the heterospecific stimulus (*P <* 0.001). Likewise, there is a difference between the number of significant inhibitory features (*P <* 0.05).

Figure 5A shows that the spike rate prediction, as measured by the scaled correlation coefficient, did not differ significantly between units stimulated with mouse USVs and units stimulated with pitch-shifted starling songs (*P* = 0.55). The relation between the measured correlation coefficient between the model prediction and the recording and the expected correlation coefficient of the associated unit is shown in Figure 5B. Raster plots for different levels of expected correlation can be seen in Figure 4. To account for any differences in firing rate between the responses from mouse USVs and pitch-shifted starling songs, the firing rates of all recorded responses in both stimulus paradigms were compared, as shown in Figure 5C. No significant difference was observed between the responses to conspecific and heterospecific vocalizations (*P* = 0.78).

**Figure 4:**
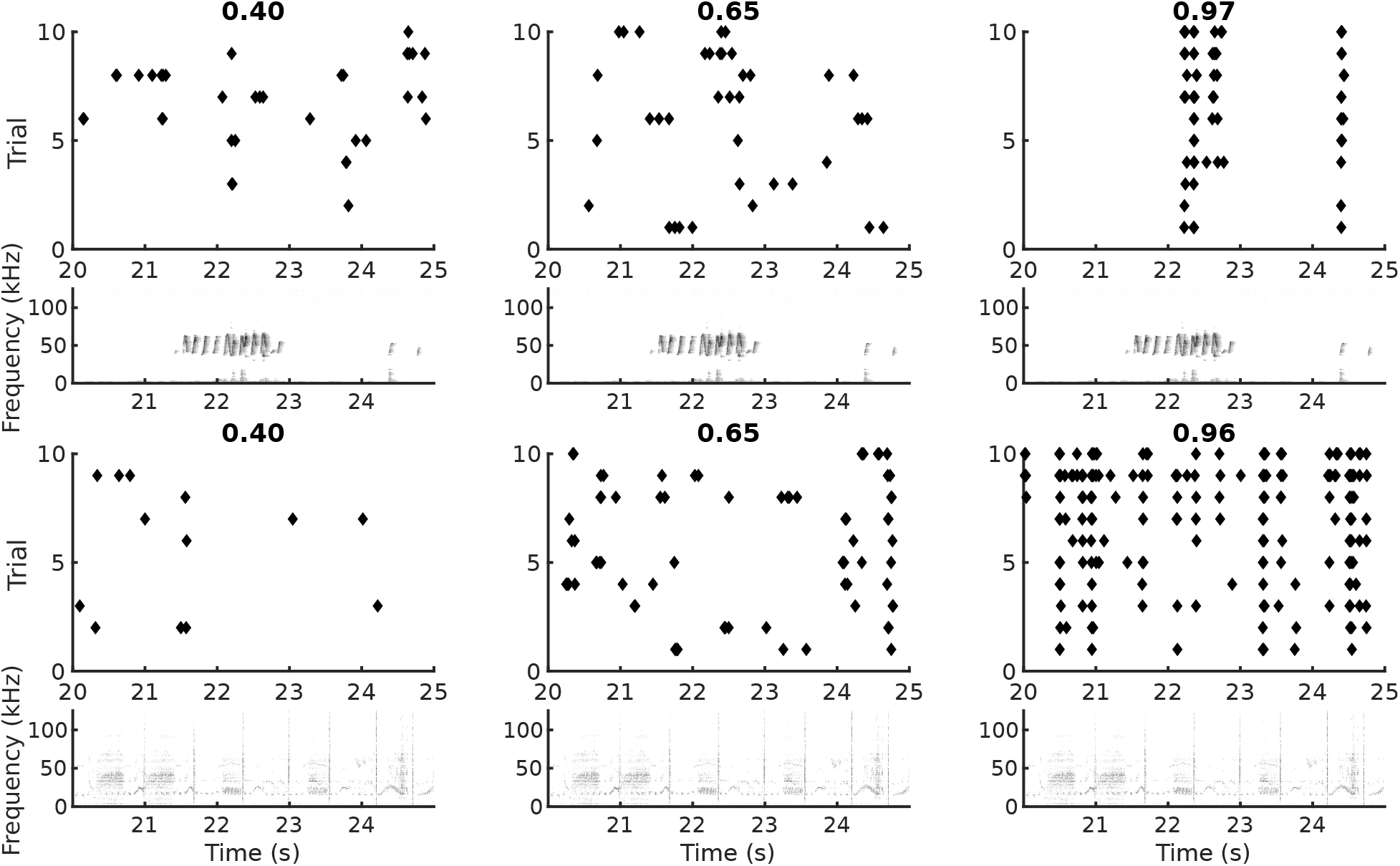
Raster plots showing three different levels of expected correlation. **(Top)** shows the expected correlation for three units responding to USVs. The expected correlation for each unit is indicated above the raster plot. The spectrogram of the associated stimulus is plotted below each of the raster plots. **(Bottom)** shows the expected correlation for three units responding to starling song. The expected correlation for each unit is indicated above the raster plot. The spectrogram of the associated stimulus is plotted below each of the raster plots. Note that the units in **(Top)** and **(Bottom)** are not the same units.

**Figure 5:**
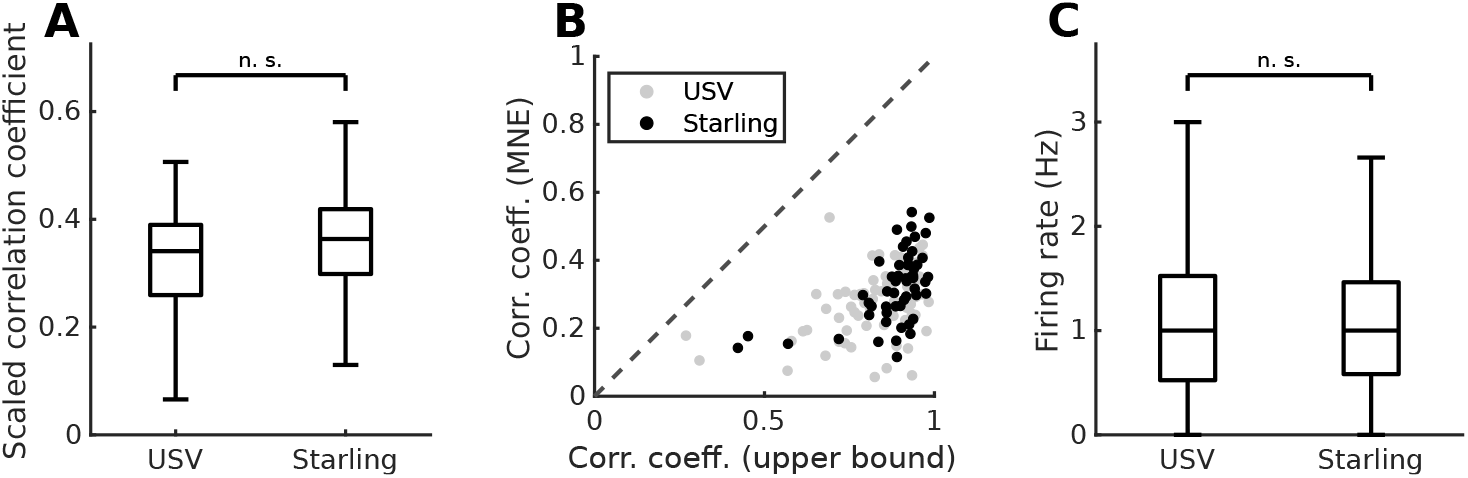
**(A)** Correlation coefficients between MNE model predictions and recorded neural activity in response to stimulation with either USVs (*n* = 74, *µ* = 0.33, *σ* = 0.11) or pitch-shifted starling songs (*n* = 54, *µ* = 0.36, *σ* = 0.10). *P* = 0.55 (two-sample Kolmogorov-Smirnov test). The correlation coefficients are scaled by the correlation coefficient between the responses to repeated presentations of the same stimulus in the test set. **(B)** Model prediction correlation coefficients plotted as a function of the expected correlation between trials to indicate an upper bound on the correlation coefficient achievable by the model. Dashed line indicates a gradient of 1. **(C)** Comparison of the firing rates of the recorded units in response to either USVs (*n* = 74, *µ* = 1.3 Hz, *σ* = 1.2 Hz) or pitch-shifted starling songs (*n* = 54, *µ* = 1.3 Hz, *σ* = 1.2 Hz). *P* = 0.78 (two-sample Kolmogorov-Smirnov test). In both cases, the firing rates were calculated using full recordings and not only test data.

### 3.3 Stimuli modelled with basis features

We calculated a set of optimal basis features for both the mouse USVs and starling songs using the sparse filtering algorithm to determine whether receptive fields are formed to represent the stimulus in a statistically optimal way. For both sets of stimuli, 128 features were calculated. Sparsity of the population activity of the features was confirmed by calculating the percentage of features with activity lower than 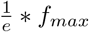, where *f*_*max*_ is the maximum activation across the entire stimulus. This was 95.2% and 92.6% for the features estimated from USVs and the starling song, respectively.

We then compared these calculated features with the estimated biological receptive fields and with the syllables from the raw stimulus from which they were calculated to determine the similarity between them. To do this, we projected them into a latent space using the UMAP algorithm (see Methods). The resulting clustering is shown in Figure 6. While the MNE features obtained with both the conspecific (USVs) and heterospecific (starling songs) stimuli formed a single cluster, only a small subset of starling syllables and USV syllables clustered together with the MNE features (1.8 % and 2.4 % respectively), the majority forming a second, separate cluster. That cluster also included the sparse-filtering features.

**Figure 6:**
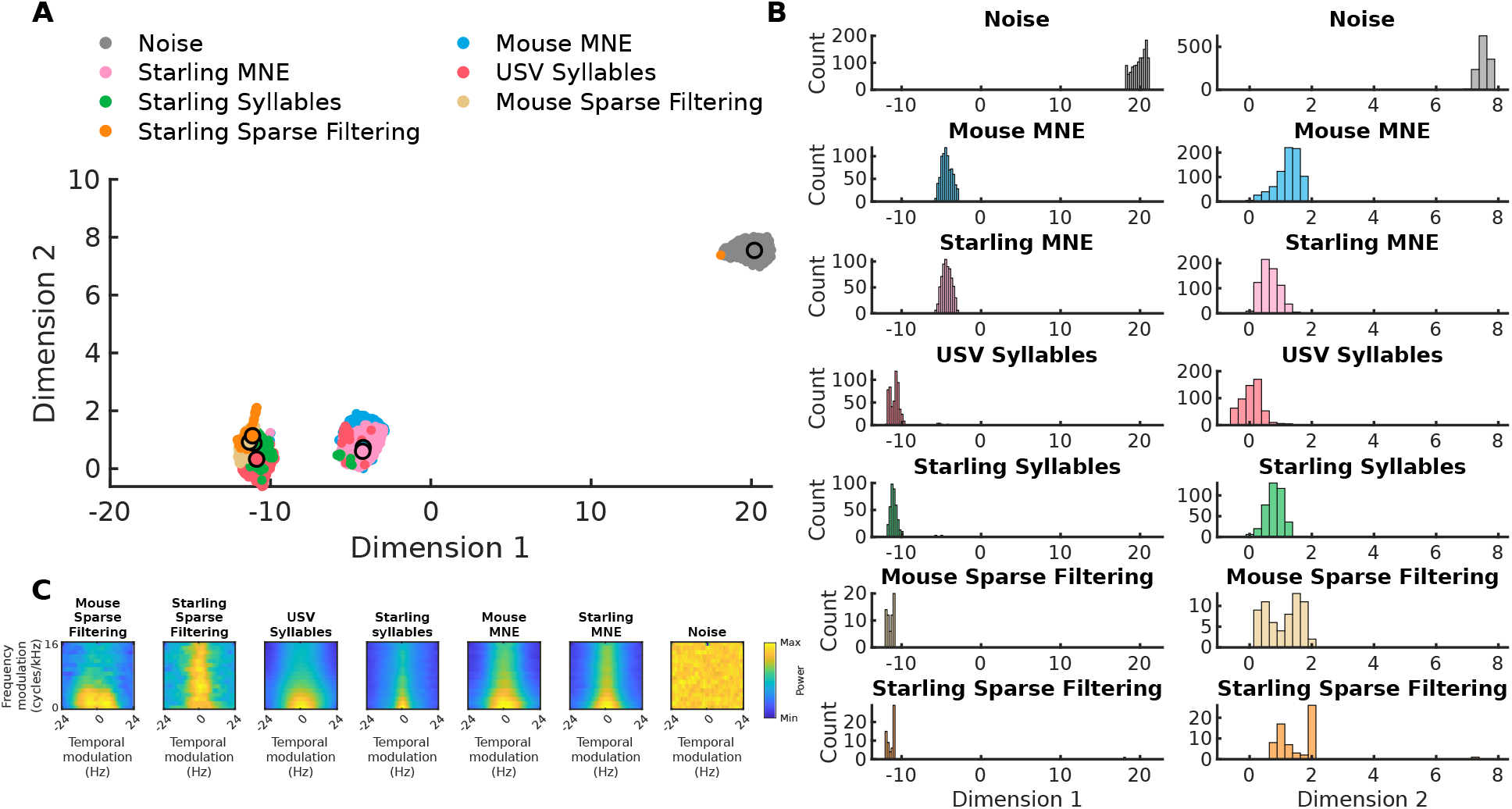
**(A)** UMAP projections onto two dimensions of: segmented syllables present in USVs and pitch-shifted starling songs, significant features estimated using the MNE models, features estimated using the sparse filtering method, and randomly generated noise. Larger points with outlines are the projection of the mean modulation power spectrum of the class as indicated by the label color. The leftmost cluster includes the USV and starling syllables segmented from the stimuli and the sparse filtering features estimated from the stimuli. The middle cluster is mainly comprised of the MNE features estimated from both stimulus types. The rightmost cluster clearly separates the noise. **(B)** Histograms of each of the different categories in dimension 1 (left) and dimension 2 (right). **(C)** Mean modulation power spectra for each of the classes in **(A)**.

## 4 Discussion

Ultrasonic vocalizations are an important communication component of social behavior in mice [Sangiamo et al., 2020]. With the ultimate aim of linking neural circuits to behavior, and given the power of the mouse as a model organism in neuroscience in general and auditory neuroscience in particular, it is useful to understand how auditory cortical circuits process these communication sounds.

To understand better the principles underlying this process, we have recorded neuronal responses (spikes) from auditory cortical neurons in mice, and characterized their receptive fields with natural stimuli. We have used mouse ultrasonic vocalizations and pitch-shifted birdsongs (as a natural but ethologically irrelevant stimulus), and employed statistical methods that work with complex natural stimuli and can identify any number of receptive-field features. Our results indicate that auditory cortical neurons in the mouse, when stimulated by USVs, display multidimensional receptive fields. This result buttresses an emerging consensus that auditory cortical neurons in birds and mammals have composite receptive fields when probed with natural complex sounds. Furthermore, the results show that a natural but ethologically irrelevant heterospecific stimulus, the pitch-shifted starling song, is an effective stimulus for the auditory cortical neurons in the mouse. Indeed, we have found that the receptive fields estimated with this stimulus have more features, both excitatory and inhibitory, than those estimated with the ethologically relevant, conspecific stimulus (USVs). However, as discussed below, the MNE features obtained with the two types of stimuli were alike, suggesting a degree of natural stimulus-invariance in the receptive fields of these neurons. The more complex starling song may simply be more efficient at sampling the stimulus space given a limited duration of experimental recordings.

Next, we have taken advantage of the developments in the field of unsupervised neural networks—implicitly or explicitly based on cortical hierarchies—which can learn statistically optimal feature representations. We have trained the sparse filtering algorithm, which was used previously to learn features of the starling song [Kozlov and Gentner, 2016], to discover statistically optimal representations of these stimuli. We have compared the biological and artificial representations using a recently developed dimensionality-reduction algorithm, UMAP, that has been previously used to capture, cluster and visualize structure of vocalizations of different animals, including songbirds and mice [Sainburg et al., 2020]. One notable outcome of this analysis is that the MNE features obtained with the conspecific and heterospecific stimuli clustered together, whereas most of the stimuli and sparse-filtering features formed a separate cluster. This implies that auditory cortical neurons in the mouse do not represent the full gamut of acoustical features present in these vocalizations. However, given the relatively low predictive power of the MNE models with these data, it would be important to investigate this question using better models of complex receptive fields, in particular models that can account for non-stationarities and capture arbitrary stimulus-response mappings [Keshishian et al., 2020].

The similar structure of auditory cortical receptive fields in the starling and in the mouse, specifically the fact that they have multiple excitatory and inhibitory subunits, and bearing in mind the ancient split between the synapsids and diapsids during amniote evolution, indicates that this property is likely a generic principle of natural sound representation.

## Conflict of interest

The authors declare no conflicts of interest.

## Acknowledgements

This work was funded by Biotechnology and Biological Sciences Research Council grant “How do auditory cortical neurons represent ethologically relevant natural stimuli? Characterizing stimulus feature selectivity and invariance” (BB/N008731/1).

## Notes

### Competing Interest Statement

The authors have declared no competing interest.

